# Coupling Raman Spectroscopy and Acoustic Microscopy to Characterize Glioblastoma Resistance

**DOI:** 10.64898/2026.10.14.759182

**Authors:** Tuna Pesen, Arda Inanc, Demet Inci, Ahmet Cingoz, Bora Akgun

## Abstract

Glioblastoma (GBM) is still the most lethal and the most treatment-resistant primary brain tumor, largely because of the cellular adaptations that let the cells escape from therapeutic stress. The genetic and the metabolic drivers of resistance are well studied, whereas the mechanochemical variations allowing GBM cells to survive inside a stiffening tumor microenvironment have received much less attention. In this study we combined Raman spectroscopy and Scanning Acoustic Microscopy (SAM) to characterize the mechanochemical signatures of drug-sensitive and drug-resistant GBM populations without any labeling. Raman spectral analysis showed different biomolecular fingerprints for the resistant cells, particularly within the protein, lipid and nucleic acid related bands, which points to a metabolic and structural remodeling. Unsupervised two-cluster classification of the spectral features recovered the biological labels with 69.5% accuracy, above the 50% chance level for two classes. The acoustic measurements showed a progressive increase in the impedance of the sensitive cells under TRAIL, while the resistant cells kept a stable and already elevated value. Oil-Red-O staining confirmed the Raman lipid-band divergence independently, the neutral-lipid content of the resistant cells being 4.1-fold greater. Resistance in this model is therefore accompanied by a distinct mechanochemical phenotype which is visible in both the biochemical and the mechanical channels. Combined vibrational and acoustic profiling may offer a practical route to identifying resistant populations at an early stage, and a biophysical basis for candidate diagnostic markers.

**SIGNIFICANCE:** Therapy resistance in glioblastoma is usually described in genetic or metabolic terms. Here we ask whether it also has a measurable biophysical form. Combining Raman spectroscopy with scanning acoustic microscopy on TRAIL-sensitive and TRAIL-resistant glioblastoma cells, we find that resistance carries both a biochemical and a mechanical signature: the resistant cells differ at protein, lipid and nucleic-acid bands, contain 4.1-fold more neutral lipid, and hold a high acoustic impedance that does not change when the drug is applied, while sensitive cells stiffen as they die. Because acoustic impedance is an absolute quantity and neither measurement requires a label, the approach offers a route to recognising resistant populations from their physical state alone.

## INTRODUCTION

Glioblastoma (GBM) is the most lethal form of primary brain cancer, with a median survival of only 12–15 months in spite of aggressive treatment regimens including surgery, radiation and chemotherapy (1). The main reason of the treatment failure is its inherent complexity together with the development of drug resistance, which lets GBM cells evade apoptosis and repopulate the tumor (2, 3). Extensive research has been carried out on the genetic mutations (4), the epigenetic modifications (5–7) and the metabolic reprogramming (8–10), however much less is known about the mechanochemical adaptations supporting therapy resistance and invasion(11).

Among the emerging targeted approaches for glioblastoma, Tumor Necrosis Factor-Related Apoptosis-Inducing Ligand (TRAIL) has attracted considerable interest, since it selectively induces apoptosis in cancer cells through the activation of death receptors (DR4 and DR5) while largely sparing normal tissues (12–14). Even though it has a promising therapeutic potential and a p53-independent mechanism of action, many GBM cells develop acquired resistance to TRAIL, which limits its clinical efficacy (15–17). The molecular basis of TRAIL resistance has been investigated extensively, but considerably less is known about the accompanying biophysical and mechanochemical adaptations that may contribute to the survival and the persistence of the resistant tumor cells.

In GBM, variations in the extracellular matrix (ECM) composition increase tissue stiffness, which acts as a critical mechanical driver of cancer progression (18–20). These mechanochemical properties, defined by the cell stiffness, the adhesion forces and cytoskeletal dynamics, are crucial in regulating how cells sense and survive therapeutic stress (21). Particulary, the plasma membrane interface undergoes a constant protein and lipid turnover in order to communicate with this stiffening surrounding (22). Consequently, the cytoskeletal remodeling at this interface is central to the adaptive response of GBM cells to the ECM variations and to the invasive behavior (19).

In spite of these observations, some questions have remained open. How does the cytoskeleton make a transition from a sensitive to a resistant phenotype? Do resistant GBM cells show specific mechanical variations promoting survival? Can these biophysical variations be used as biomarkers for the early detection of resistance? Answering such questions requires advanced label-free biophysical tools which are able to probe both the biochemical and the mechanical properties.

Raman spectroscopy and Scanning Acoustic Microscopy (SAM) come forward as promising techniques for the study of diseases including brain cancer (23–30). Raman spectroscopy provides molecular fingerprints and detects biochemical variations such as protein conformation and lipid composition changes. In parallel, acoustic microscopy works on a different contrast, mapping the intracellular elasticity and stiffness non-invasively and at high resolution, and it therefore reports on the cytoskeletal reinforcement accompanying therapeutic evasion. The two techniques have been applied to brain tumor samples separately, but to our knowledge not together on the same sensitive/resistant pair. Molecular biology techniques give a snapshot of the genetic expression and frequently miss the interplay between the physical state of the plasma membrane and the underlying metabolic shifts. In the present study SAM and Raman spectroscopy were applied to TRAIL-sensitive (A172-S) and TRAIL-resistant (A172-R) sublines, in order to record the mechanical and the biomolecular fingerprints of the same phenotypes, and to see whether the biophysical variations can serve as candidate markers of the resistant state.

The structural integrity and biochemical composition of the glioblastoma microenvironment are intrinsically linked, where mechanical stiffness often dictates the cellular signaling pathways associated with chemoresistance. Although traditional molecular biology techniques provide a snapshot of genetic expression, they often overlook the dynamic interplay between the physical state of the plasma membrane and the underlying metabolic shifts. Accordingly, in the present study we combined SAM with Raman spectroscopy on TRAIL-sensitive (A172-S) and TRAIL-resistant (A172-R) sublines, in order to record the mechanical and the biomolecular fingerprints of the same phenotypes, and to see whether the biophysical variations can be used as candidate markers of the resistant state.

## MATERIALS AND METHODS

### Cell Culture Preparation

Human glioblastoma A172 cell lines were obtained from the American Type Culture Collection (ATCC, USA). TRAIL-sensitive (A172-S) and TRAIL-resistant (A172-R) sublines were previously generated by Cingöz et al. (31), in which parental A172 cells were rendered resistant to TRAIL through prolonged exposure. These established sublines were utilized in the present study. Cells were cultured in Dulbecco’s Modified Eagle Medium (DMEM; Gibco, USA) supplemented with 10% fetal bovine serum (FBS; Gibco, USA) and 1% penicillin–streptomycin (Capricorn, Germany). All cells were maintained in a humidified incubator at 37°C with 5% CO_2_. The culture medium was replaced every 2–3 days, and cells were passaged upon reaching appropriate confluency using standard cell culture procedures. TRAIL was either purchased commercially (SuperKiller, Enzo Life Sciences, USA) or produced in HEK293T cells as previously described (32).

### Cell Viability Assays

Cell viability was evaluated using the Cell Titer-Glo Luminescent Cell Viability Assay (Promega, USA). A172-S and A172-R cells were seeded in black, clear-bottom 96-well plates (Corning Costar) at a density of 104 cells/well and incubated overnight to allow for attachment. Cells were treated the following day with TRAIL (5-10-25 ng/ml), with each condition performed in triplicate. After 24 hours of treatment, an equal volume of Cell Titer-Glo reagent was added to each well. Plates were incubated at room temperature to induce cell lysis and stabilize the luminescent signal. Luminescence was recorded using a Cytation 5 Cell Imaging Multi-mode Reader (BioTek Instruments, USA). All experimental values were normalized to the mean of the untreated control group and the result of the viability test is given in Fig. 1.

**Figure 1:**
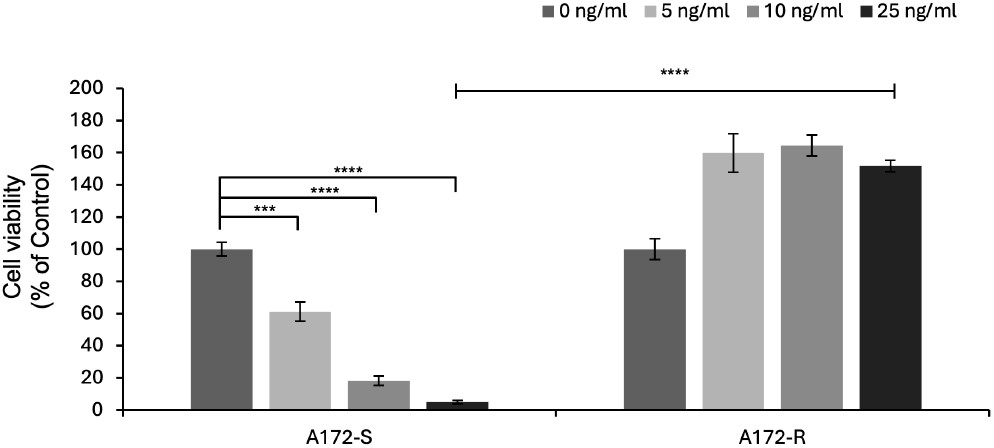
Viability of TRAIL-sensitive and TRAIL-resistant A172 cells in response to 24-h treatment of different TRAIL doses (0-25 ng/ml). (*** and **** denote p < 0.05, p < 0.001 respectively, two-tailed Student’s t-test).

### Sample Preparation for Acoustic Microscopy

Both drug-sensitive (A172-S) and drug-resistant (A172-R) glioblastoma cells were cultured 24 hours prior to the experiments. For acoustic impedance mapping, A172-S and A172-R cells were seeded into 35-mm specialized culture dishes at a density of 3 × 105 cells per dish in 1.5 mL of complete growth medium. To minimize acoustic attenuation and ensure high-resolution signal acquisition, Petri dishes with an ultra-thin bottom thickness of 50 *μ*m were utilized. The samples were incubated overnight under standard conditions (37°C, 5% *CO*_2_) to ensure optimal adherence and physiological stability. Following incubation, cells were treated with TRAIL at the specified concentrations, while untreated cells served as negative controls. Acoustic impedance measurements were acquired 3–4 hours post-treatment using a high-frequency Scanning Acoustic Microscopy (SAM) system (AMS-50SI, Honda Electronics, Toyohashi, Japan) using 320 MHz transducer. Samples were measured after 4 hours of TRAIL treatment and subsequently recorded at 45-minute intervals (Group 1 and Group 2) to assess temporal changes.

### Acoustic Microscopy

To ensure high-fidelity signal transmission and minimize acoustic energy loss, the cells were imaged within their culture medium using Petri dishes with an ultra-thin bottom thickness of 50 *μ*m. A transducer with a 320 MHz central frequency was operated in acoustic impedance mode to capture the mechanical properties of the cells, and ultrasound gel was used to couple acoustic wave to the Petri dish through the cells as illustrated in Fig. 2.

**Figure 2:**
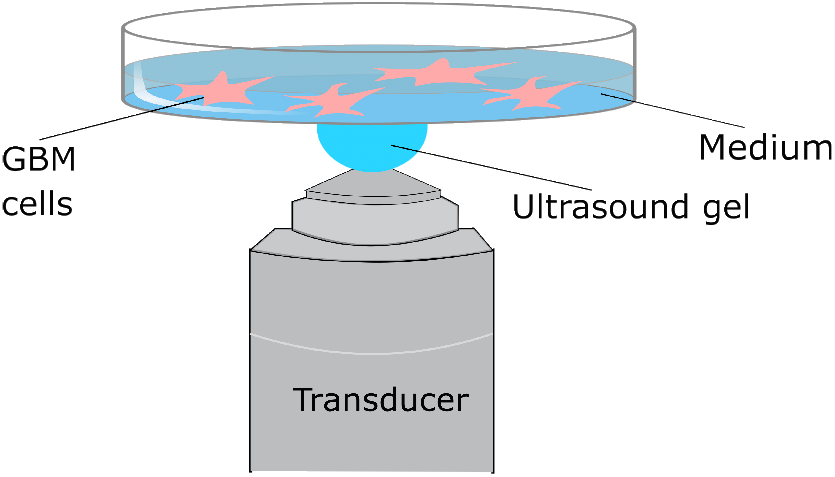
Acoustic impedance measurement of GBM cells.

For each sample, a scanning area of 0.3 mm × 0.3 mm was mapped with an acquisition time of approximately 2 minutes per frame. The data matrix was imbinarized using the Otsu’s edge detection method so as not to include the region without the cells. The steps are shown in the supplementary Fig. S1. After the edge detection of the cells, the distributions of the two cell types were box plotted.

### Sample Preparation for Raman Spectroscopy

For Raman spectroscopic analysis, A172-S and A172-R cells were seeded onto specialized stainless steel slides, to ensure a background-free optical substrate, 24 hours prior to measurements. Upon reaching the desired confluency, the cells were fixed using cold methanol to preserve the biomolecular integrity of the intracellular components. Following fixation, the slides were meticulously rinsed with deionized water to remove residual media constituents and crystalline salts that could interfere with the spectral signal. The samples were then allowed to air-dry for 15 minutes.

### Raman Spectroscopy

In the experimental setup, an optical microscope with 40× objective lens (Zeiss EC Epiplan NA = 0.60), a Raman spectrometer (Ocean QE Pro, Orlando FL, USA), and a continuous laser source with a wavelength of 785 nm and a power of 500 mW (Laser Glow, Toronto, Canada) were utilized (Fig. 3). The collimated laser beam was directed to the sample using a silver mirror to pump the ground vibrational state of the molecules in GBM cells. The Raman scattered photons coming from the sample (the dark red beam in Fig. 3) were passed through the dichroic mirror and directed to the spectrometer through a lens (f=50 mm) and two sequential Raman edge filters (Thorlabs FELH0800), which were used to eliminate the Rayleigh scattered photons. The spectrometer collected the photons through a multimode fiber with a numerical aperture (NA) of 0.22. The samples were scanned using a motorized XY scanning stage (Standa 8MTF-102LS05), which was linked to MATLAB.

**Figure 3:**
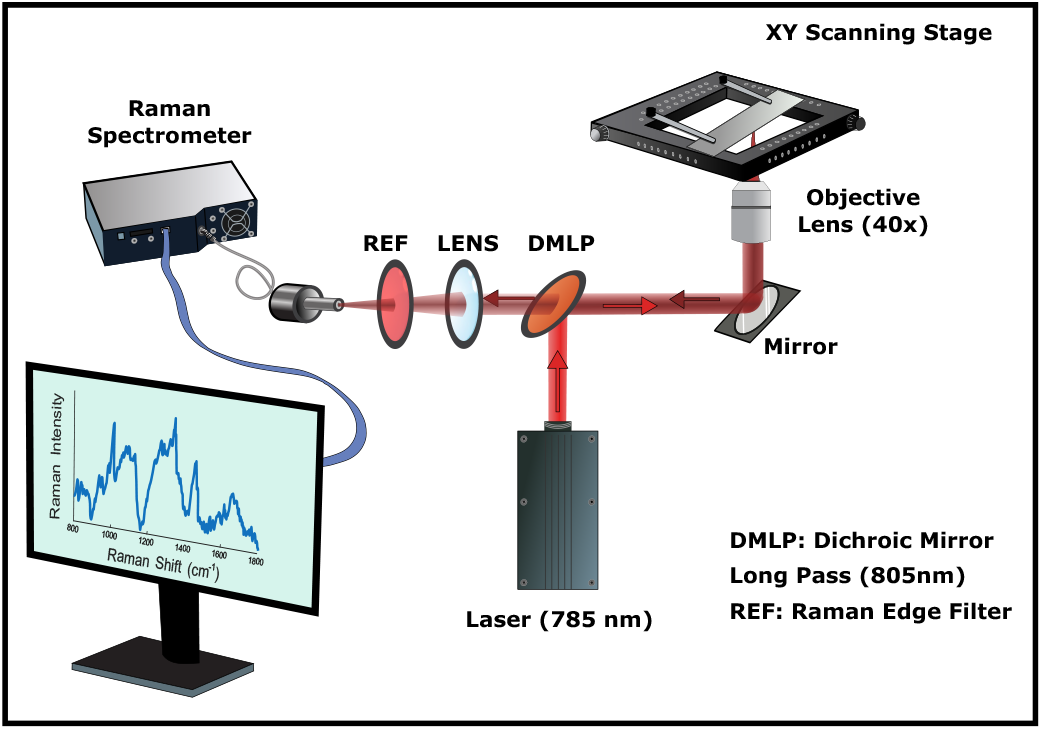
Raman spectroscopy setup.

The pre-processing of the Raman spectral data includes calibration, baseline subtraction, and vector normalization. To perform fluorescence baseline removal, a fifth order polynomial spline fit curve was subtracted from each spectrum. Finally, a vector normalization method (also called L2 normalization) was applied to the spectra for better comparison.

The pixels of the metal region around the cells were disregarded using Principal Component Analysis (PCA) and OPTICS clustering algorithms. Thus, only the spectra from the cells were included in the further analysis.

For the analysis of the spectra, the mean spectra are compared (standard deviations along with the mean spectra are given in the supplementary Fig. S3). Additionally, a one way analysis of variance (ANOVA) was conducted and the most discriminative peaks were determined using effect size (*η*^2^ calculation). The dimensional reduction algorithm t-SNE was applied before clustering. After the dimensions of the data were reduced to two, it was clustered using the OPTICS algorithm. All these steps were performed on MATLAB R2025a.

### Oil-Red-O Lipid Quantification

Neutral-lipid content of the cells was quantified from Oil-Red-O–stained brightfield images of TRAIL-resistant (A172-R) and TRAIL-sensitive (A172-S) cells (four fields per group) as shown in Fig. 4.

**Figure 4:**
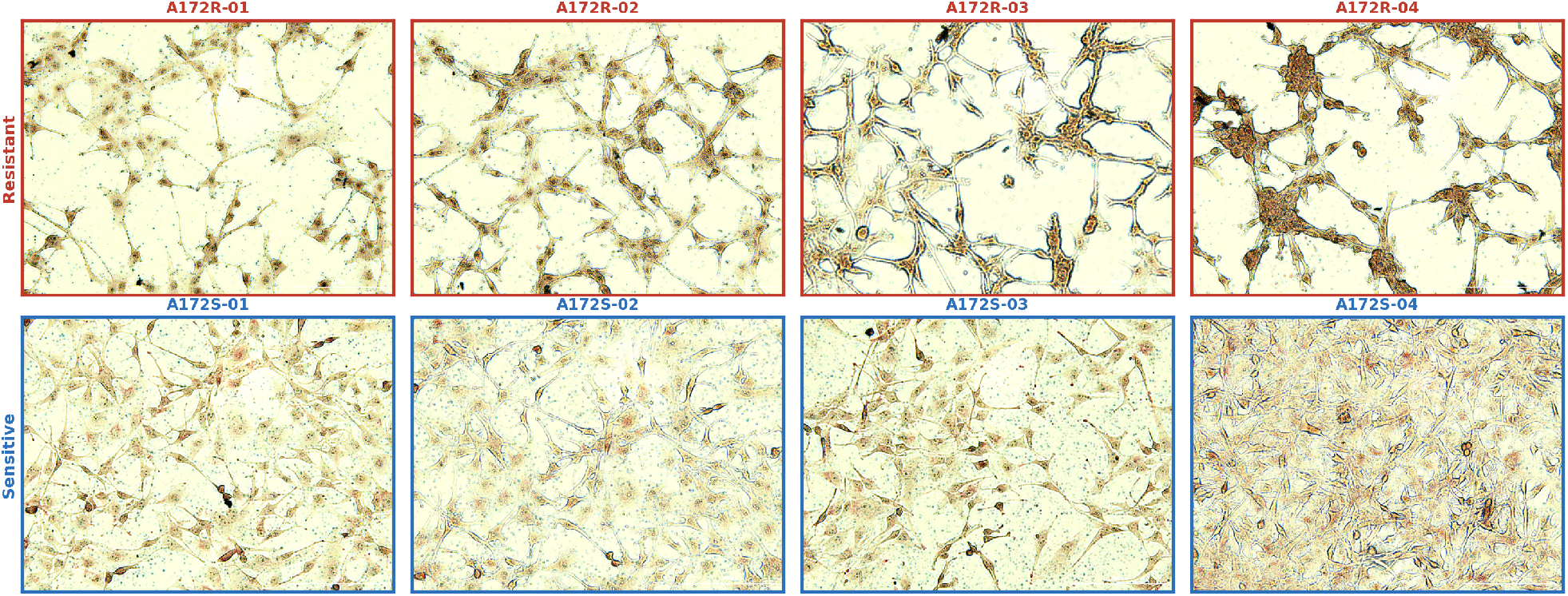
Representative Oil-Red-O–stained brightfield fields of resistant and sensitive A172 glioblastoma cells used for lipid quantification. Neutral lipids were visualized by Oil-Red-O staining and imaged under brightfield microscopy at 10× magnification. Top row (red): TRAIL-resistant A172-R fields; bottom row (blue): TRAIL-sensitive A172-S fields (n = 4 fields per group). Resistant cells form a sparse, branched network with prominent perinuclear Oil-Red-O accumulation, whereas sensitive cells display a more homogeneous, evenly spread distribution with weaker staining.

Cell-covered regions were delineated by semi-automated segmentation in MATLAB, and all subsequent analysis was restricted to the intracellular area (Fig. 5).

**Figure 5:**
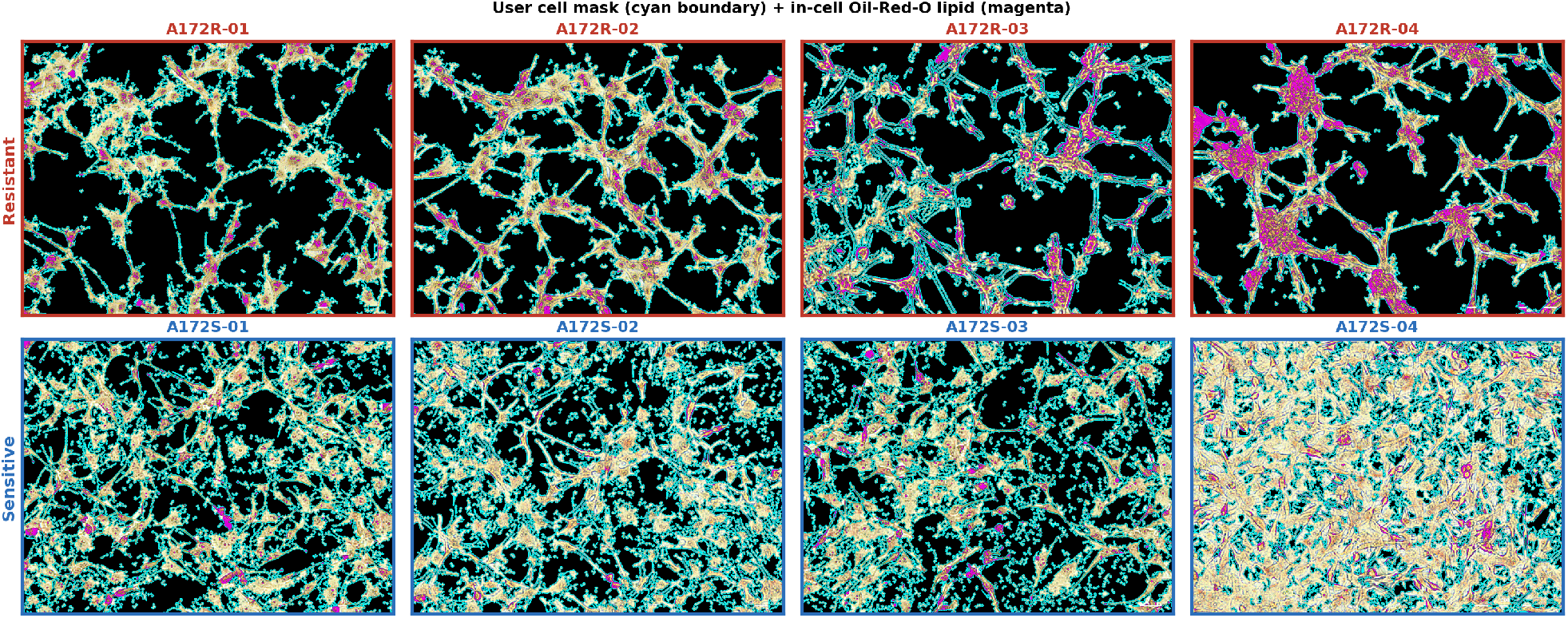
MATLAB segmentation for Oil-Red-O quantification. Cell masks (cyan boundary) with in-cell lipid-positive pixels (magenta) for four resistant and four sensitive 10× fields, using a common global Otsu threshold.

Within each cell mask, images were white-balanced to their in-cell bright (unstained) reference and converted to optical density, and the Oil-Red-O signal was isolated by colour deconvolution. A single global intensity threshold (Otsu’s method computed on the pooled in-cell stain signal of both groups) was applied identically to every field to define lipid-positive pixels, ensuring an unbiased comparison. For each field we computed the lipid-positive fraction of cell area, the mean in-cell stain optical density, and the integrated optical density per cell pixel. Group means were compared at the field level using a two-sided Welch t-test and a Mann–Whitney U test, with Cohen’s d as the effect size.

## RESULTS

### Acoustic Impedance Measurement Results

According to Fig. 6, the TRAIL-sensitive and TRAIL-resistant GBM cells exhibited pronounced changes in their acoustic impedance. To quantify the time-dependent cellular response to TRAIL, an edge-detection algorithm was applied to the acoustic impedance data. (See Fig. S1 and S2) The measurements were calibrated by assigning the background medium as water with an acoustic impedance of 1.49 MN·s/m3 (1.49 MRayl). The acoustic impedance difference of the sensitive and resistant cells measured at 45-min intervals is shown in Fig. 7. Under the effect of TRAIL, the acoustic impedance of the sensitive cells was found to increase progressively over time. In contrast, the TRAIL-resistant cell group showed no appreciable temporal variation in acoustic impedance following TRAIL treatment.

**Figure 6:**
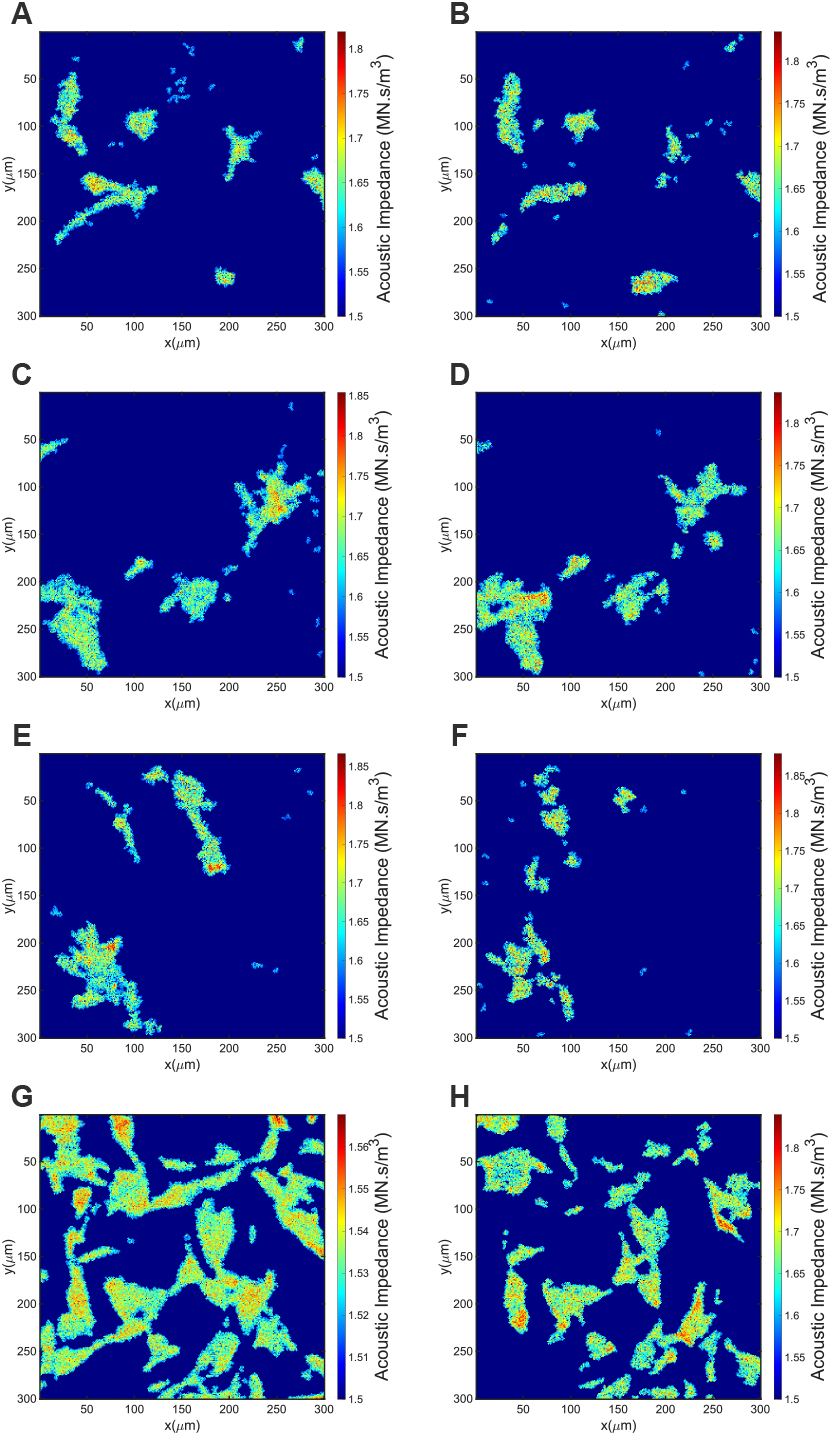
Time-resolved acoustic impedance maps of resistant and sensitive GBM cells under TRAIL treatment. The dynamic mechanical response was captured at two distinct time points with a 45-minute interval. A,C,E) Resistant GBM cells after 4 hours following TRAIL treatment (*t* = 0). B,D,F) The same resistant cells after 45 minutes of exposure, demonstrating mechanical resilience. G) Sensitive GBM cells at *t* = 0 and H) sensitive cells after 45 minutes, showing characteristic impedance shifts. The color scale represents quantitative acoustic impedance values in MN·s/m3 (MRayl).

**Figure 7:**
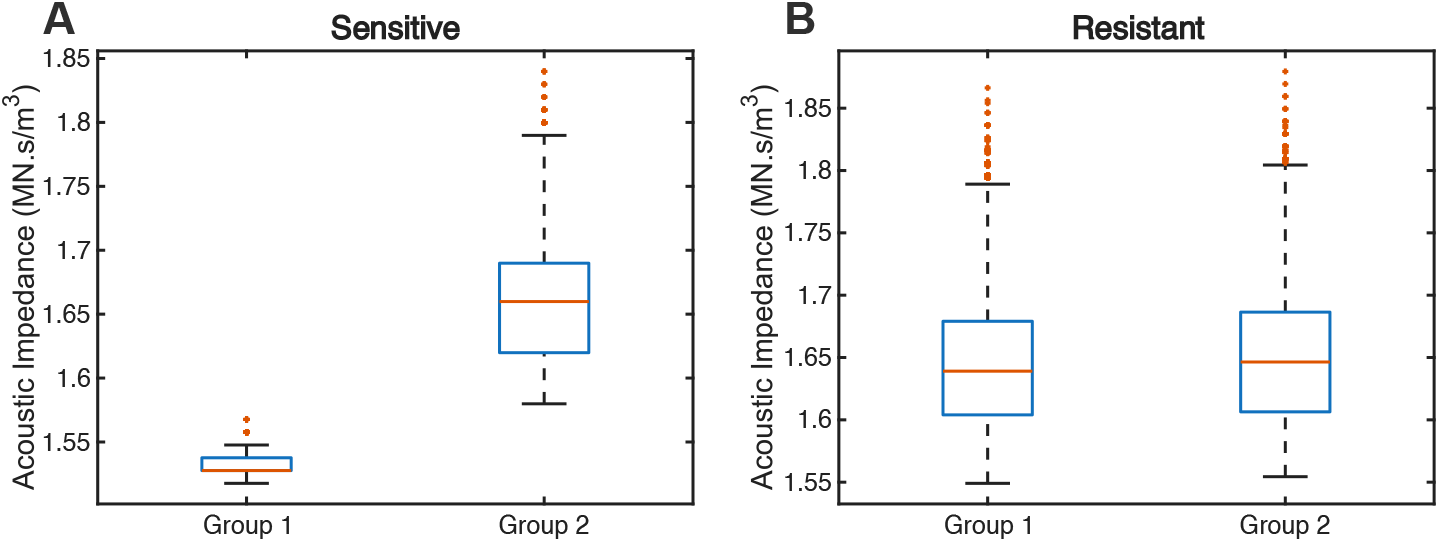
a) Acoustic impedance response of sensitive glioblastoma cells to TRAIL with 45 minute apart (Group 1 median = 1.50 MN·s/m3, Group 2 median = 1.66 MN·s/m3), b) Acoustic impedance response of resistant glioblastoma cells to TRAIL with 45 minute apart (Group 1 median = 1.64 MN·s/m3, Group 2 median = 1.65 MN·s/m3).

### Raman Spectral Measurement Results

The resistant and sensitive groups comprised 980 and 904 Raman spectra, respectively. The mean Raman spectra of the sensitive and resistant groups are presented in Fig. 8a. The main discriminative four Raman bands are determined through calculated *η*^2^ values with a common threshold of 0.14 as shown in Fig. 8b (33). The four most discriminative Raman bands are highlighted in cyan in Fig. 8a.

**Figure 8:**
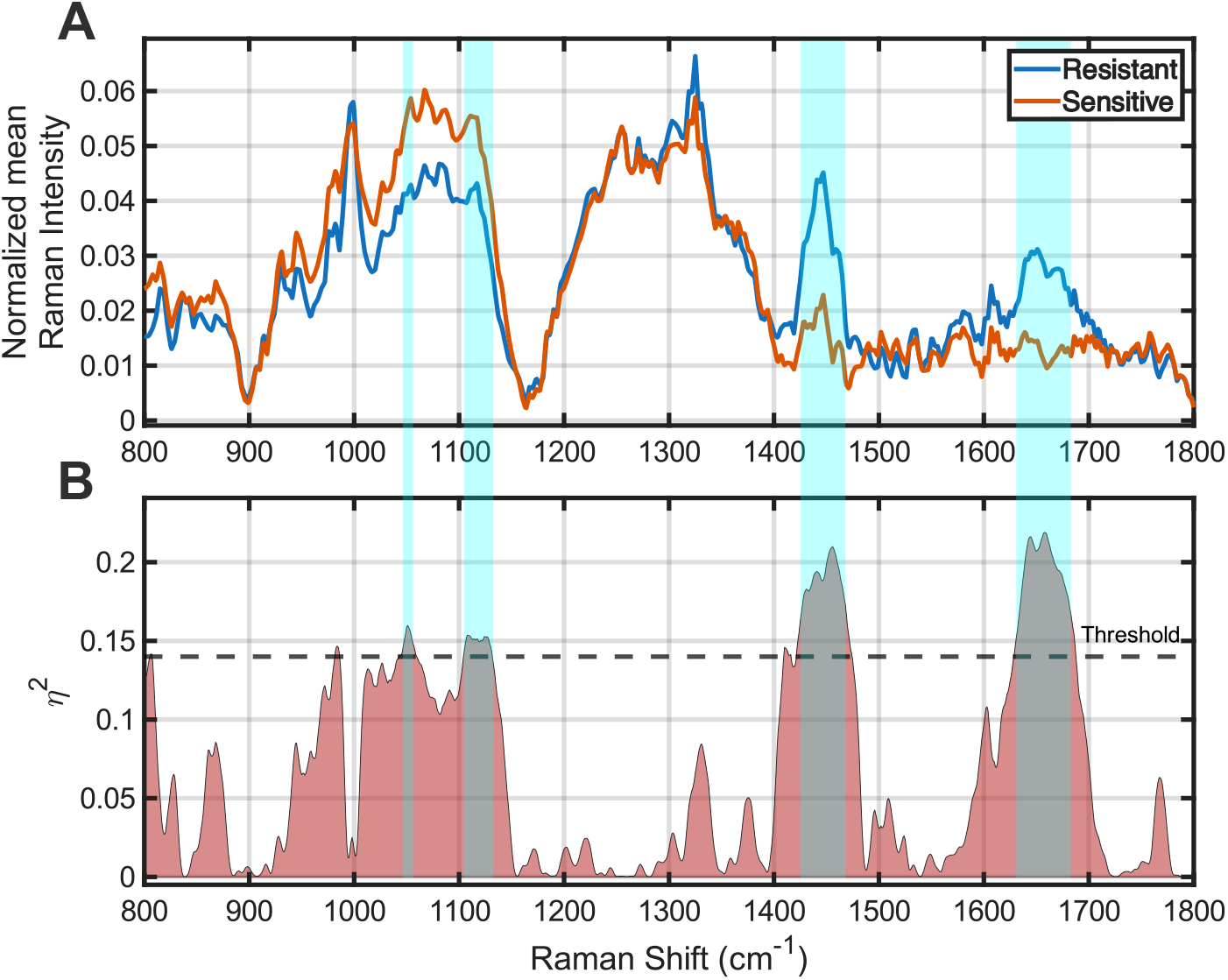
a) Normalized mean Raman intensity of resistant and sensitive GBM cells. b) Calculated *η*^2^ values for ANOVA effect size. The most prominent differences on four regions are highlighted in cyan on both panels, namely around the peaks at 1658 cm−1, 1456 cm−1, 1108 cm−1, 1051 cm−1.

The corresponding p-values found to be p < 0.001 for the maximum peaks at these regions and the peaks at 1658 cm−1, 1456 cm−1, 1108 cm−1 and 1051 cm−1 were attributed predominantly to protein related contributions (the Amide I band), lipid associated vibrations (such as CH_2_ bending and C-C stretching), phospholipid and nucleic acid signatures (PO_2_− modes), and C-C stretching of lipids, respectively. Furthermore, the rather pronounced separation observed within the 1400–1700 cm−1 spectral region appears to be indicative of alterations, mainly in terms of membrane lipid composition and accompanying changes in protein secondary structure, which, taken together, seem to constitute the dominant biochemical distinctions between the two groups. Relative to the sensitive cells, the resistant spectra were higher at 1658 and 1456 cm−1 and lower at 1108 and 1051 cm−1. Since the spectra were vector normalized, these differences describe how the spectral weight is distributed between the two windows rather than absolute concentrations.

Unsupervised two-cluster classification of the Raman spectra is given in (Fig. 9a), with the corresponding ground-truth labels in Fig. 9b.

**Figure 9:**
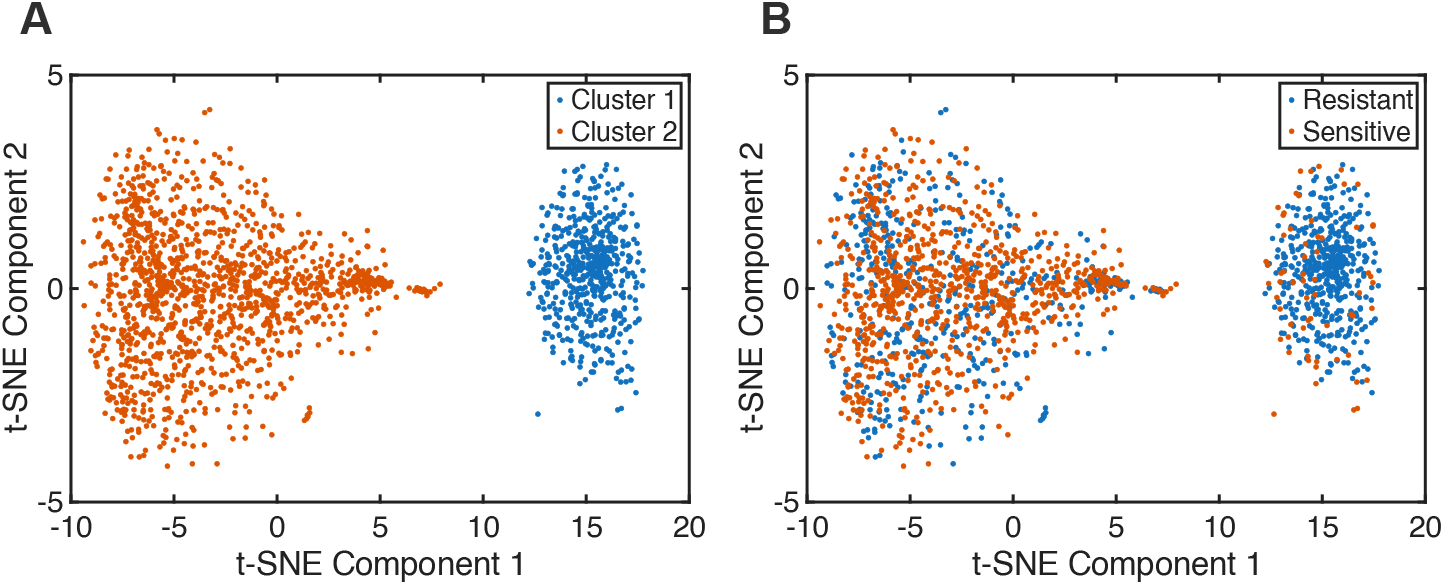
Unsupervised clustering analysis of Raman spectra. a) Cluster assignments based on spectral similarities without prior labeling. b) Ground truth labels indicating drug-sensitive (orange) and drug-resistant (blue) GBM cells. The model demonstrates a clear spatial separation primarily along Component 1, reflecting intrinsic biomolecular differences.

The clustering reached an overall accuracy of 69.5%, so the spectra organize into two partially separable populations even without the class labels. In the confusion matrix the two error rates are not symmetric: 469 of the 980 resistant spectra were assigned to the resistant cluster (47.9%), whereas 841 of the 904 sensitive spectra were assigned correctly (93.0%). The overall accuracy is therefore carried mainly by the sensitive class. The overlap region is therefore populated mainly by resistant spectra that fall on the sensitive side, which is what would be expected if the resistant population were the more heterogeneous of the two rather than if both phenotypes contributed equally (Fig. 10).

**Figure 10:**
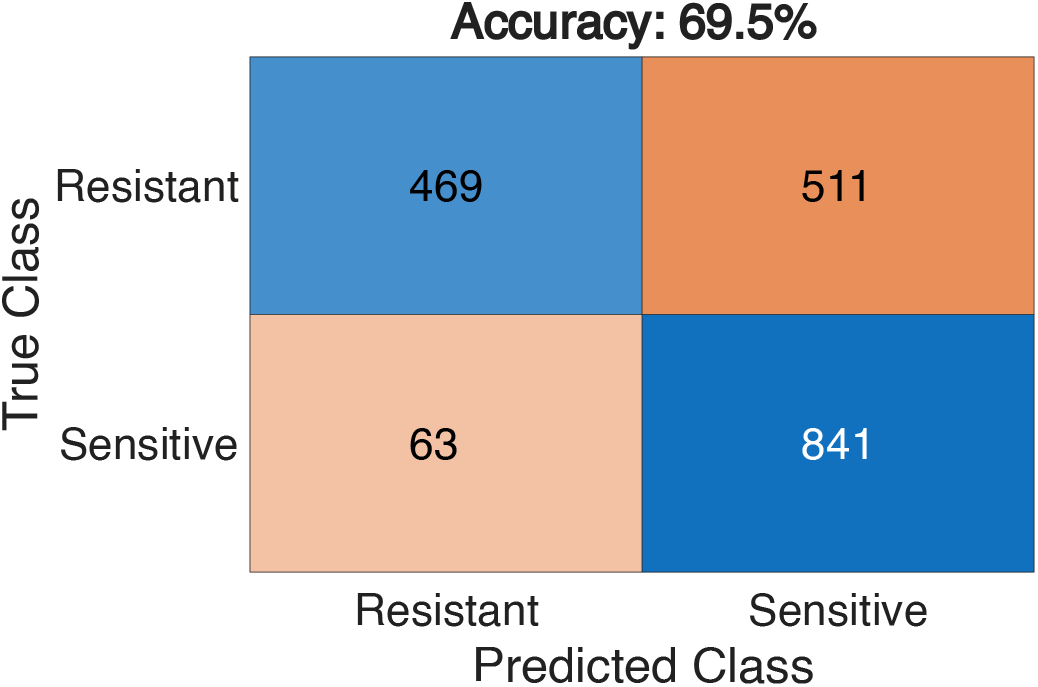
Confusion matrix and accuracy of unsupervised classification.

### Oil-Red-O Quantification Results

Oil-Red-O quantification revealed markedly higher neutral-lipid content in resistant cells, consistent with the Raman lipid-band divergence (1456 cm−1) as given in Fig. 11. When normalized to cell area, the lipid-positive fraction was 4.1-fold greater in A172-R than in A172-S (12.4 ± 7.5% vs 3.1 ± 1.6%; mean ± s.d., n = 4 fields per group), and every resistant field exceeded every sensitive field (two-sided Mann–Whitney U, p = 0.029). The same direction was seen in the mean in-cell stain density (0.43 vs 0.25 OD) and the integrated optical density per cell pixel, and the effect size was large (Cohen’s d = 1.74; Welch t-test, p = 0.084, the parametric test being conservative here because one resistant field was particularly lipid-rich and inflated the within-group variance). Thus, both the per-area lipid coverage and the staining intensity indicate greater intracellular lipid accumulation in resistant cells, providing an independent, image-based confirmation of the stronger CH_2_ band at 1456 cm−1 in the Raman spectra.

**Figure 11:**
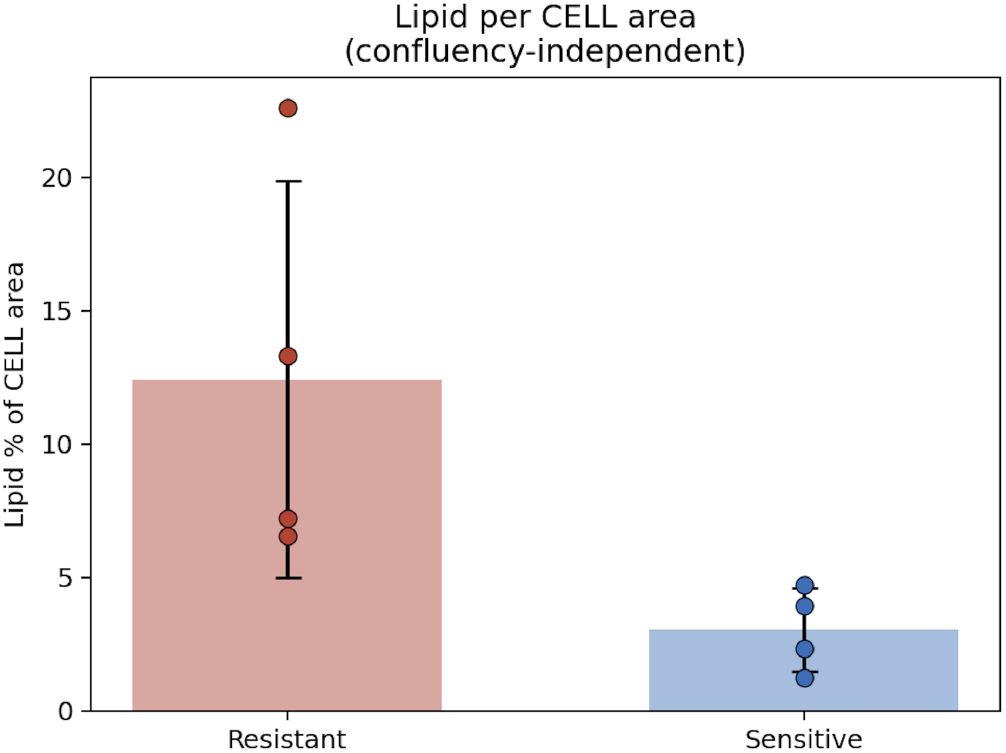
Oil-Red-O–positive area as a percentage of cell area in resistant (red) and sensitive (blue) A172 cells. Bars, mean; error bars, s.d.; points, individual fields (n = 4/group). Resistant cells show 4.1-fold higher intracellular lipid (Mann–Whitney U, p = 0.029; Cohen’s d = 1.74).

## DISCUSSION

The 1000–1150 cm−1 region carries the vibrational signatures of the phosphate backbone of phospholipids and nucleic acids, together with C–O and C–C skeletal modes. In the A172-R population the intensities in this window, and in particular the 1108 cm−1 band, were lower than in the sensitive cells, while the CH_2_ deformation band at 1456 cm−1 was higher. Taken together these two observations describe a change of composition rather than a change confined to one class of molecule: neutral lipids such as triacylglycerols and cholesteryl esters are rich in CH_2_ groups but carry no phosphate head group, so a cell that stores more of them gains weight at 1456 cm−1 and loses it between 1000 cm−1 and 1150 cm−1. The vector normalization applied to the spectra makes this redistribution explicit.

Resistant GBM cells are also known to change their nucleic acid and mitochondrial content. Kaur et al. showed that radiation-resistant cells survive lethal stress by forming multinucleated giant cells through homotypic fusion, and that the resulting populations carry an increased genomic content (34), while Wang et al. reported enlarged mitochondria and an increased mitochondrial mass in TMZ-resistant GBM cells (35) and Nakhle et al. showed that mitochondrial acquisition from mesenchymal stem cells rewires the metabolism towards nucleotide biosynthesis (36). All of these would add signal in the 1000–1150 cm−1 window, and our spectra move in the other direction. The nucleic acid and mitochondrial pools are therefore not what drives the separation in this model, and the band cannot in any case distinguish a nuclear from a mitochondrial contribution. What the two windows do follow is the lipid content.

The clustering was unsupervised, and t-SNE preserves the local neighborhoods rather than maximizing the class separation. These two partially distinct clusters therefore reflect an intrinsic difference in the biochemical organization of the spectra, and not a separation engineered with prior label knowledge. An accuracy of 69.5% is modest, so the incomplete separation obtained here is more probably related with the real heterogeneity of the two GBM populations than with the pipeline itself. GBM is heterogeneous and the transition between the two states is gradual rather than binary. Furthermore, the *η*^2^ analysis pointed to the same lipid, protein and nucleic acid bands, which suggests that the individual Raman shifts combine into a multidimensional fingerprint. The shifts in the lipid and protein regions may also indicate a mesenchymal-like transition (37), frequently reported for recurrent and therapy-resistant glioblastoma (38, 39).

Lipid remodeling is the part of the picture with the most independent support. Choo et al. reported altered cell morphology in TMZ-resistant GBM cells, together with increased cholesterol and fatty acid synthesis, reduced lipid unsaturation and decreased membrane fluidity (40). More recently, Naghibzadeh et al. identified a stable lipid-resistant state with lysophospholipid enrichment, sphingolipid accumulation, cholesteryl ester storage and extensive vesicular remodeling (41). Our Oil-Red-O measurements go in the same direction: the lipid-positive fraction of the cell area was 4.1-fold greater in the A172-R cells, and every resistant field exceeded every sensitive field. The cholesteryl ester storage described by Naghibzadeh et al. is of this kind. Wang et al. examined resected GBM tissue with label-free Raman spectromicroscopy and showed that the neutral lipid held in these droplets is mainly triacylglycerol and cholesteryl ester, and that the stored amount tracks the MGMT methylation status (42). Stored neutral lipid of this kind raises the CH_2_ band without adding to the phosphate region, which is the pattern we observe. The lower intensity at 1051 cm−1 may also carry a contribution from the carbohydrate C–O modes that fall there, which would fit a shift towards lipid-based metabolism. Chakraborty et al. described a lipid-dependent lineage of treatment-resistant persister cells which establish metabolically specialized interactions with the tumor-associated macrophages in order to secure lipid resources (43). Therefore, lipid accumulation may not be only a consequence of resistance.

The acoustic data given in Fig. 7 add a mechanical dimension to these variations. The resistant cells started from a greater impedance than the sensitive ones, the medians being 1.64 MN·s/m3 and 1.50 MN·s/m3, and did not move under TRAIL, the median after 45 minutes being 1.65 MN·s/m3. In contrast, the sensitive cells stiffened from 1.50 MN·s/m3 to 1.66 MN·s/m3, and apoptotic condensation is the straightforward explanation for this second observation. What distinguishes the resistant phenotype is then not a high stiffness in itself, but a high baseline which does not respond to the drug. Such a behaviour would follow from a denser cytoskeleton and an altered plasma membrane, and this kind of shielding may limit the drug penetration and help the cells to withstand the stiffening extracellular matrix. The impedance is a product of density and sound speed, however, and says nothing directly about the viscoelasticity.

Multinucleation, mitochondrial expansion, metabolic rewiring, membrane remodeling and a mechanical adaptation have all been reported separately for resistant GBM. The Raman and the acoustic measurements reported here are compatible with that set, and both were obtained without any label. The agreement between these resistance-associated properties and our spectroscopic and acoustic observations suggests that label-free Raman and acoustic analysis may be used to detect resistant states, and to follow the mechanochemical processes underlying glioblastoma therapy resistance.

## LIMITATIONS

Several limitations should be stated. The work rests on a single cell line pair, A172-S and A172-R. The acoustic part samples two time points only, 45 minutes apart, so the temporal behaviour is coarse. The 69.5% figure comes from the Raman spectra alone; the two modalities were recorded on separate samples and were never combined in a single classifier. Moreover, the Raman measurements were made on methanol-fixed cells and the acoustic measurements on live cells in their medium, and this difference in the sample state should be kept in mind when the two are compared. One further point concerns the impedance itself, which in the reflection mode is affected by the local cell thickness as well as by the material properties, so a part of the contrast reported here may be morphological rather than intrinsic. A quantitative height measurement, for example by atomic force microscopy or quantitative phase imaging, would separate the two contributions. A wider panel of GBM lines, more time points and a joint classifier are the obvious next steps.

## CONCLUSION

Drug resistance in the A172 glioblastoma model is accompanied by a mechanochemical phenotype which can be detected without labels. The Raman measurements separated the two populations at the protein, lipid and phosphate bands (1658, 1456, 1108 and 1051 cm−1), the resistant cells being stronger at 1658 and 1456 cm−1 and weaker at 1108 and 1051 cm−1. This is the pattern expected when a cell stores neutral lipid, which is rich in CH_2_ groups and free of phosphate head groups. Furthermore, the Oil-Red-O quantification supported the lipid part independently, the neutral-lipid content being 4.1-fold greater in the resistant cells.

Scanning acoustic microscopy added the mechanical side of the same comparison: at the first time point the resistant cells already carried a greater acoustic impedance than the sensitive ones (medians 1.64 against 1.50 MN·s/m3), and this value did not move over the following 45 minutes under TRAIL (1.65 MN·s/m3), whereas the impedance of the sensitive cells rose progressively (to 1.66 MN·s/m3).

For the early detection of therapeutic resistance this is a starting point and not a finished tool. The measurements are label-free and they need little sample preparation. Such a combination is what makes the approach worth developing further for treatment monitoring in glioblastoma management.

## AUTHOR CONTRIBUTIONS

T.P., A.C., A.I. and B.A. designed the research, D.I. prepared the samples. T.P., A.I. and D.I. analyzed the data. T.P., A.I., D.I., A.C., and B.A. wrote the manuscript. All the authors reviewed the manuscript.

### ACKNOWLEDGMENTS

This study was supported by the Presidency of the Republic of Türkiye, Presidency of Strategy and Budget (SBB) (Grant No. 2026K12-283013).

## SUPPLEMENTARY MATERIAL

**Figure S1:**
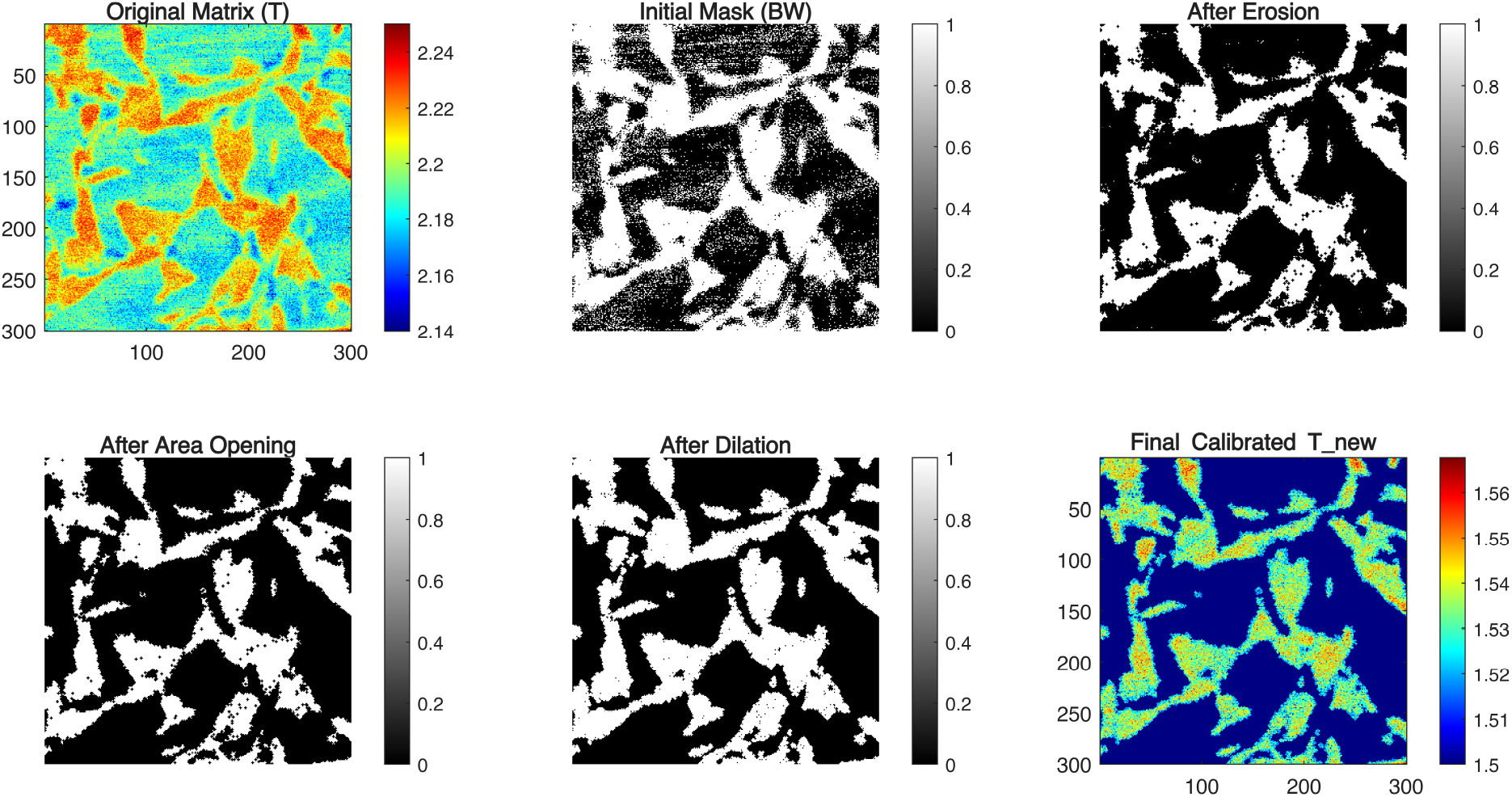
Edge detection process for sensitive Group 1 cells.

**Figure S2:**
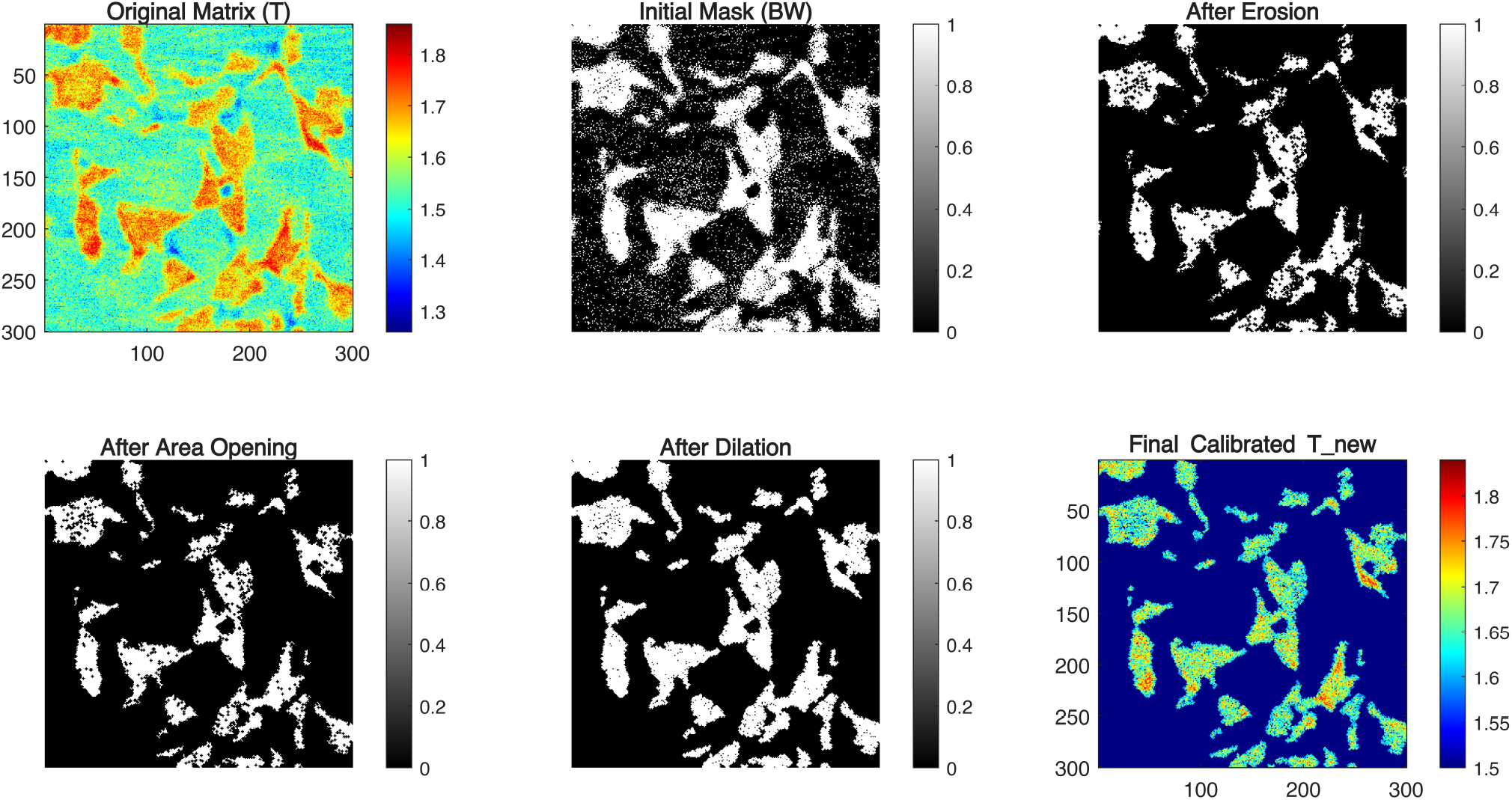
Edge detection process for sensitive Group 2 cells.

**Figure S3:**
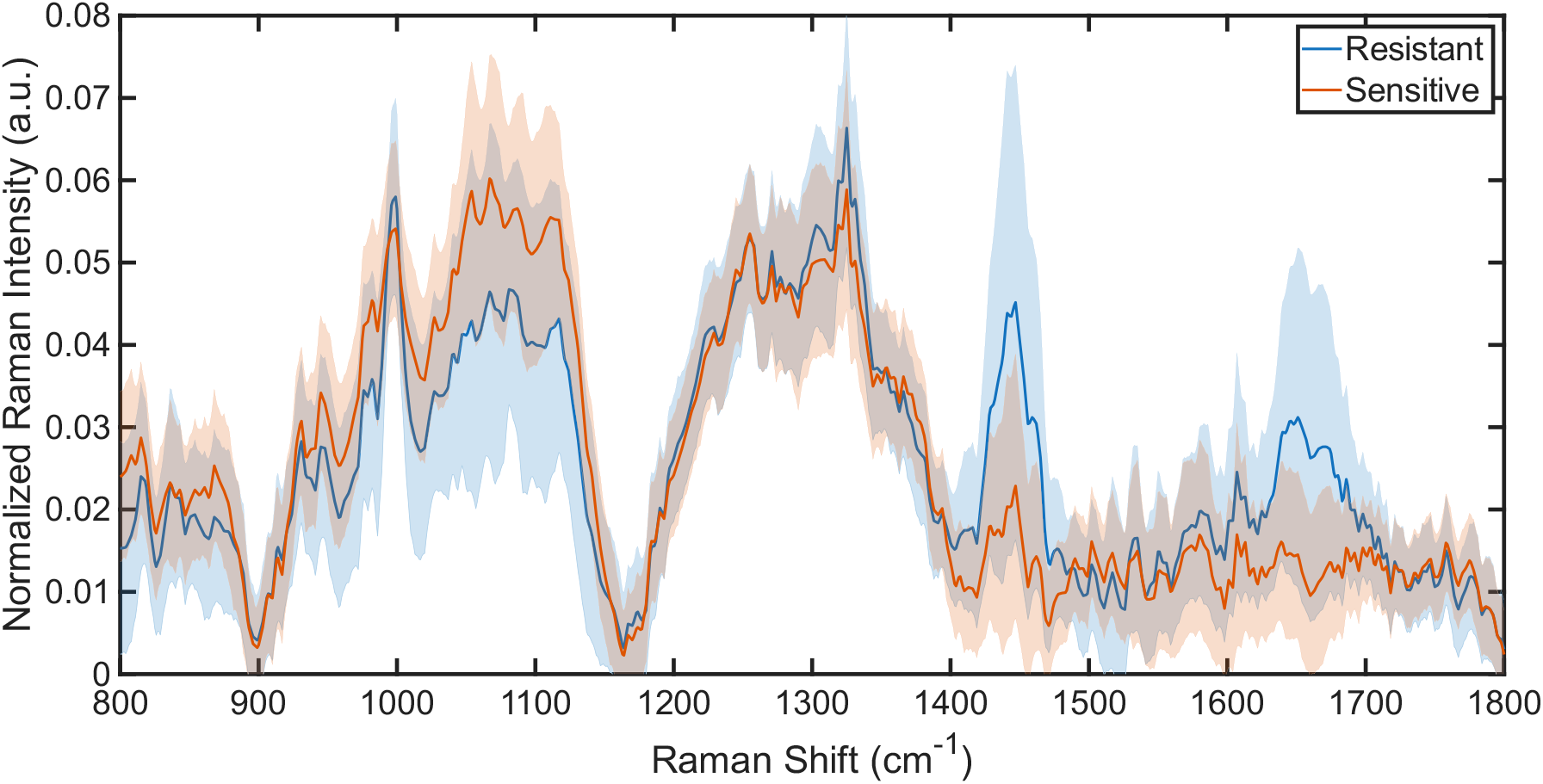
Normalized mean Raman intensity with standard deviation of resistant and sensitive GBM cells.

